# The Hydroxyproline O-arabinosyltransferase *FIN4* is required for tomato pollen intine development

**DOI:** 10.1101/2022.08.16.504199

**Authors:** Syeda Roop Fatima Jaffri, Holly Scheer, Cora A. MacAlister

## Abstract

The pollen grain cell wall is a highly specialized structure composed of distinct layers formed through complex developmental pathways. The production of the innermost intine layer, composed of cellulose, pectin and other polymers, is particularly poorly understood. Here we demonstrate an important and specific role for the hydroxyproline O-arabinosyltransferase (HPAT) *FIN4* in tomato intine development. HPATs are plant-specific enzymes which initiate glycosylation of certain cell wall structural proteins and signaling peptides. *FIN4* was expressed throughout pollen development in both the developing pollen and surrounding tapetal cells. A *fin4* mutant with a partial deletion of the catalytic domain displayed significantly reduced male fertility *in vivo* and compromised pollen hydration and germination *in vitro.* However, *fin4* pollen that successfully germinated formed morphologically normal pollen tubes with the same growth rate as the wild-type pollen. When we examined mature *fin4* pollen, we found they were cytologically normal, and formed morphologically normal exine, but produced significantly thinner intine. During intine deposition at the late stages of pollen development we found *fin4* pollen had altered polymer deposition, including reduced cellulose and increased detection of pectin, specifically homogalacturonan with both low and high degrees of methylesterification. Therefore, *FIN4* plays an important role in intine formation and, in turn pollen hydration and germination and the process of intine formation involves dynamic changes in the developing pollen cell wall.

## Introduction

One of the key innovations of land plant evolution was the development of pollen as a method for transporting and protecting male gametes. While the motile sperm of early land plants required free liquid water through which the sperm could swim, pollen are resistant to desiccation and often released from the anther in which they develop in a partially dehydrated form (Hackenberg and Twell, 2019). Upon contact with a compatible pistil, the pollen will hydrate, increasing in volume and germinate, forming a pollen tube. This pollen tube grows through the maternal tissue of the flower until it reaches a receptive ovule where it will ultimately rupture, releasing its sperm nuclei cargo and allowing fertilization to proceed (Dresselhaus and Franklin-Tong, 2013). This process is essential for successful flowering plant sexual reproduction and many important crops, including tomato, require fertilization to trigger seed production and fruit set (Giorno et al., 2013).

The cell wall of the mature pollen grain has two major layers with distinct compositions, developmental origins and biological functions (Heslop-Harrison, 1968). The protective outer layer or exine is composed of sporopollenin containing mainly long-chain fatty acids, phenylpropanoids, phenolics, and traces of carotenoids (Hess & Frosch, 1994; Jiang et al., 2013; Piffanelli et al., 1997; Souza et al., 2009). Due to its complex and heterogeneous structure, sporopollenin is resistant to most enzymes and is one of the most resilient biopolymers (Grienenberger and Quilichini, 2021). The inner layer or intine has a composition similar to other plant cell walls and is largely composed of cellulose, pectins including homogalacturonan (HG, β-1,4-galacturonic acid) and cell wall-associated proteins (Fang et al., 2008; Hess, 1993; Persson et al., 2007; Suárez-Cervera et al., 2002). The pollen tube is initially formed as an extension of the intine emerging from an aperture (i.e. a region of the pollen grain lacking exine) and is largely composed of pectin (Cascallares et al., 2020).

The formation of the pollen wall takes place in the anther during the late stages of pollen development and requires participation of both the developing haploid pollen and the surrounding diploid tapetal cell layer (Shi et al., 2015). Since the intine is largely covered by the chemically inert exine, access for biochemical studies and visual screening for intine defects is difficult, complicating the study of the molecular pathways of intine formation. Though several mutants have been described with exine phenotypes, few intine-specific mutants have been reported and those that have been described often also result in general pollen degeneration and late stage pollen abortion phenotypes (Blackmore et al., 2007; Cankar et al., 2014; Dobritsa et al., 2011; Jiang et al., 2013; Mi et al., 2022; Moon et al., 2013; Quilichini et al., 2015; Schnurr et al., 2006; Shi et al., 2015; Shim et al., 2022; Ueda et al., 2013; Van Damme et al., 2006; Wang et al., 2022). Several of these known intine mutants highlight specific cell wall polymers and glycosylation as important to proper intine development. For example, triple mutants of *Arabidopsis thaliana* cellulose synthase complex members *cesa6, cesa9,* and *cesa2* have abnormally thick intines and pollen fertility defects (Persson et al., 2007). Pectin remodeling mutants also display intine phenotypes. HG, the primary form of pectin, is initially synthesized with a high abundance of methyl-ester groups and is enzymatically de-esterified by cell wall-resident pectin methylesterases (PMEs). Removal of the methyl-ester groups allows the formation of calcium salt bridges between neighboring polymers, rigidifying the HG and strengthening the cell wall (Levesque-Tremblay et al., 2015). Arabidopsis *pme48* pollen produce intine layers with a high degree of methyl-esterified HG (meHG) leading to pollen hydration and germination defects (Leroux et al., 2015). Similarly, knockdown of several pectin degrading enzymes (polygalacturonases and pectate lyases) has been shown to impact intine formation in *Brassica campestris* pollen (Huang et al., 2009; Jiang et al., 2014a; Jiang et al., 2014b; Lyu et al., 2015). Rhamnogalacturonan I (RGI) is another major cell wall pectic polysaccharide. Arabinan RGI side chains have been implicated in the regulation of desiccation tolerance of cell walls (Moore et al., 2008). Potato with reduced pectic arabinan content show disrupted intine formation, eventually leading to loss of the cytoplasm through the thin intine at the aperture (Cankar et al., 2014). In addition to RGI, arabinose is incorporated into several other classes of cell wall polymers and glycoproteins (Kotake et al., 2016). Multiple lines of evidence support an important role for arabinose, rather than a specific arabinose-containing molecule, in pollen development and intine formation. UDP-arabinopyranose mutases, also known as reversibly glycosylated polypeptides (RGPs), convert the more energetically favorable UDP-L-arabinopyranose (UDP-Ara*p*) into the arabinofuranose form (UDP-Ara*f*) which is used by many glycosyltransferases to incorporate *Araf* into various polysaccharides and glycoproteins (Rautengarten et al., 2011). Arabidopsis *rgp1 rgp2* double mutants fail to produce a well-defined intine before arresting late in pollen development (Drakakaki et al., 2006). A similar phenotype results from knockdown of rice UDP-Arabinopyranose Mutase3 (Sumiyoshi e t al., 2015). Furthermore, rice mutants of the L-arabinokinase-like gene, *collapsed abnormal pollen1*, are also male sterile and largely missing intine (Ueda et al., 2013).

Ara*f* is covalently linked to hydroxyproline (Hyp) residues in certain protein sequence contexts by Hydroxyproline O-arabinosyltransferases (HPATs), glycosyltransferase (GT) 95 enzymes which catalyze the addition of a single β-L-Ara*f* sugar (Shpak et al., 1999; Ogawa-Ohnishi et al., 2013). A second and third β-L-Ara*f* can also be added by members of the GT77 family and a fourth, α-linked Ara*f* may be added by extensin arabinose deficient (ExAD), a GT47 family member (Egelund et al., 2007; Gille et al., 2009; Møller et al., 2017; Velasquez et al., 2011). The resulting short linear arabinose chain (Hyp-Ara) promotes an extended, rod-like protein conformation (Shinohara and Matsubayashi 2013; Stafstrom & Staehelin, 1986; Van Holst and Varner, 1984). The EXTENSINs (EXTs) are a large family of cell wall structural glycoproteins that are heavily modified in this way (Lamport and Miller, 1971; Petersen et al., 2021). Following glycosylation, EXTs are secreted into the cell wall space where they are able to form intra and inter-molecular covalent crosslinks through tyrosine residues (Held et al., 2004). This EXT network is hypothesized to function as a scaffold for pectin assembly, forming extensin pectate (Cannon et al., 2008; Lamport et al., 2011). In addition to the heavily Hyp-Ara modified EXTs, other targets of this modification include EXT chimeras and secreted signaling peptides of the CLE (CLAVATA3/Embryo Surrounding Region-Related) family (Lara-Mondragón et al., 2022; Ohyama et al., 2009; Petersen et al., 2021). The *HPAT* family is strongly conserved across plant genomes and in the absence of HPAT activity, no detectable Hyp-Ara is formed (MacAlister et al., 2016).

Here we report the function of a previously uncharacterized *HPAT, FIN4,* which promotes the formation of the intine in tomato *(Solanum lycopersicum*) pollen. The tomato genome encodes four putative *HPATs,* named *FINs* after the fasciated inflorescence phenotype of the first tomato member of this gene family characterized (Xu et al., 2015). The fasciation of *fin* mutants traces to a loss of Hyp-Ara glycosylation of CLE peptides which act as suppressors of shoot meristem cell division (Xu et al., 2015). Other reported *hpat* loss-of-function phenotypes suggest altered cell wall properties, likely due to disrupted glycosylation of EXT and/or EXT-like proteins. In the model moss *Physcomitrium patens*, *hpata* single mutants and *hpata/hpatb* double mutants have increased biomass due to faster elongation of tip-growing vegetative cells, an effect correlated with cell wall gene expression changes and which can be suppressed by addition of exogenous cellulose to the growth media (MacAlister et al., 2016). In

*Arabidopsis,* double mutants of *hpat1* and *hpat2* display root hair elongation defects (Velasquez et al., 2015) and thinner hypocotyl cell walls (Ogawa-Ohnishi et al., 2013). Arabidopsis *HPAT1* and *HPAT3* are redundantly required for pollen tube elongation and male fertility (MacAlister et al., 2016). The *hpat1 hpat3* double mutant pollen tubes have reduced rates of elongation and are prone to rupture. Our observations on *FIN4’s* specific role in the formation of the intine thus provide insight into the role of Hyp-Ara in the production of this specialized cell wall.

## Materials and methods

### Sequence alignment

The genomic sequences of *FIN3* (*Solyc07g021170*) and *FIN4* (*Solyc12g044760*) were retrieved from the Solanaceae Genomics Network (Fernandez-Pozo et al., 2015). Alignment of genomic sequences was done using JDotter (Brodie et al., 2004). HPAT protein sequences were aligned by CLUSTALW.

### Plant material and growth conditions

Tomato plants *(Solanum lycopersicum*) c.v. MicroTom (Carvalho et al., 2011; Meissner et al., 1997) were grown in trays or pots in soil, under 16-hour light: 8-hour dark cycles, in temperature-controlled growth chambers, maintained at 23°C. The *fin4* mutant was generated through a CRISPR/Cas9-based gene targeting method for simultaneous knockout of *FIN3* and *FIN4* as previously described (Brooks et al., 2014). We introgressed the *fin4* mutation from the original M82 cultivar into MicroTom over three generations and confirmed by PCR that both the CRISPR/Cas9 transgene and the *fin3* mutation were lost before phenotypic analysis. Due to the high sequence similarity between *FIN3* and *FIN4* (93.8% sequence identity), a multi-stage genotyping assay was designed to distinguish between wild-type (WT) and mutant alleles of the two genes. In the first stage, high fidelity primers (table 1) were used with high annealing temperatures (62°C for *FIN3* and 67°C for *FIN4*) to amplify a long fragment (2404 base pairs (bp) for *FIN3* and 2419 bp for *FIN4*) followed by a short PCR to distinguish between the WT and deletion alleles. For the initial long PCR, less than 5 ng of Qiagen DNeasy Plant kit (catalog number 69104)-purified genomic DNA was amplified for 15 cycles in 10μl reactions with Phusion Taq polymerase (New England Biolabs, catalog number M0530S). The long reaction products were diluted 1 to 10, and 1μl of the diluted product was used as a template in the corresponding short reaction, amplified for 25 cycles and separated on a 2% agarose gel. FIN3S primers produced 316 and 166 bp products for WT and *fin3,* respectively while the FIN4S primers amplified 291 and 229 bp fragments for WT and *fin4,* respectively.

**Table 1.**
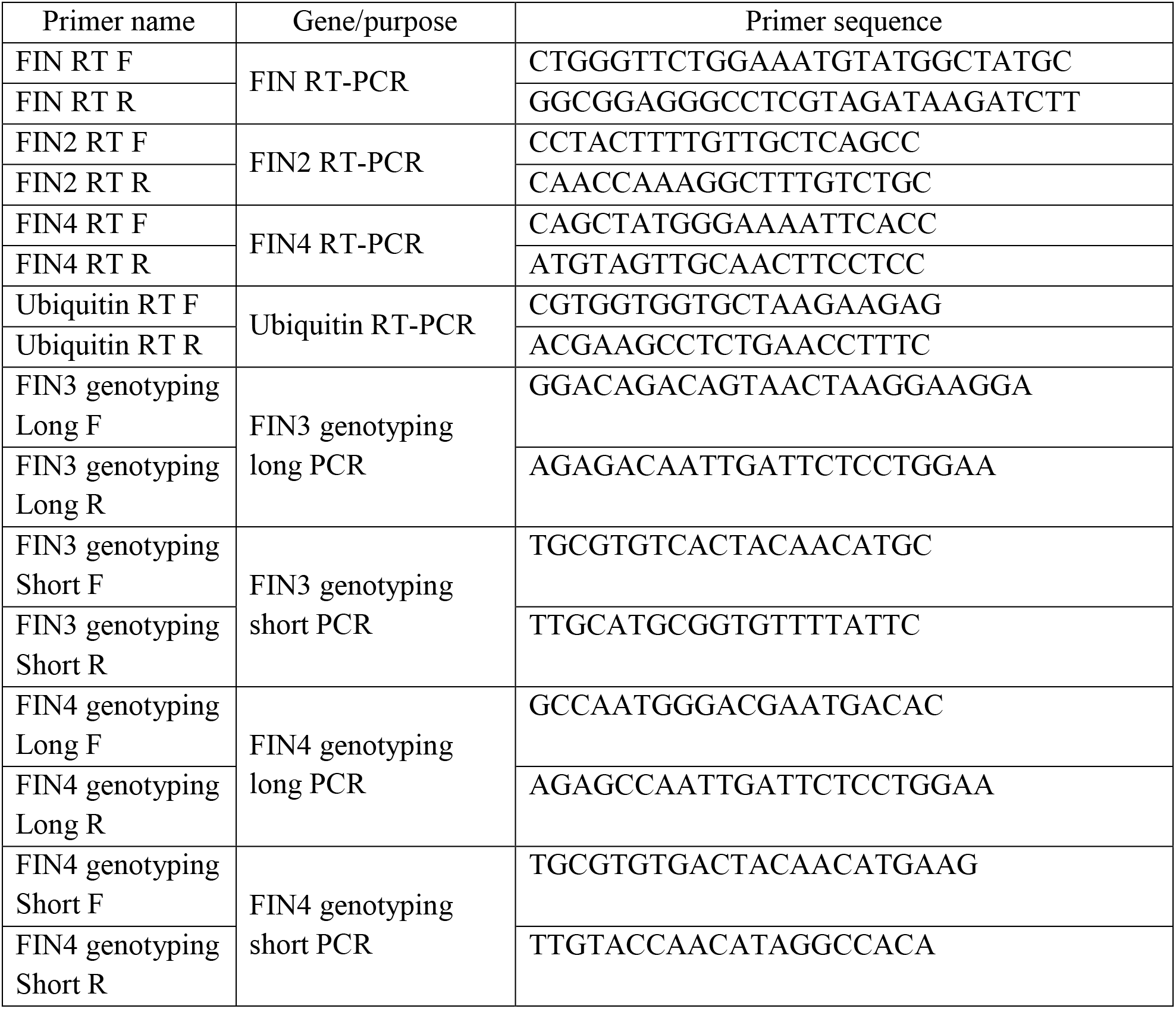
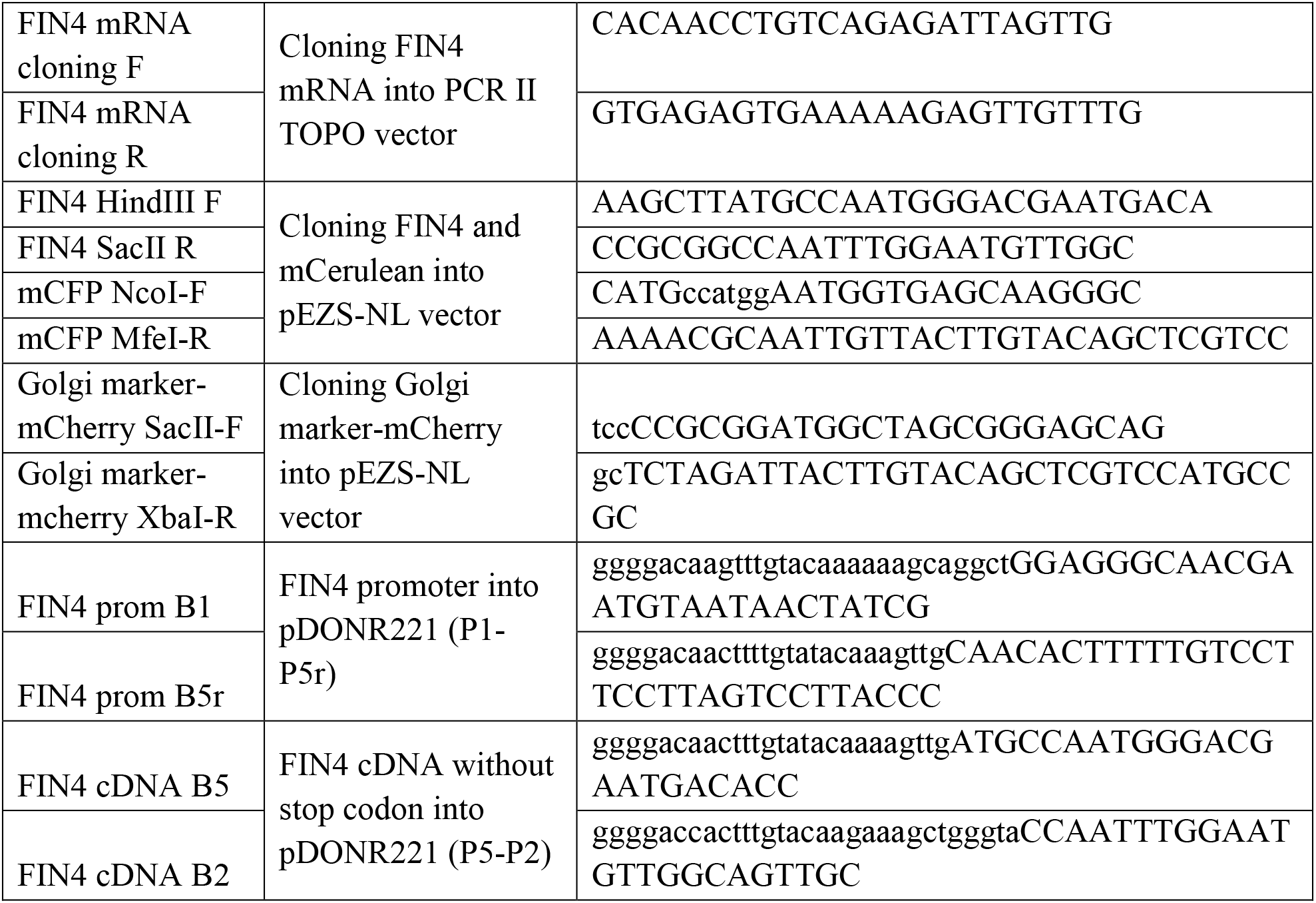
List of primers used.

### RT-PCR

Harvested tissue of the relevant organ was frozen in liquid nitrogen and then lysed by Qiagen Tissue Lyser II (catalog number 85300). Total RNA was extracted from 100 mg of tissue using RNeasy Mini Kit (Qiagen, catalog number 74004). 1 μg of extracted RNA was converted to cDNA using Superscript IV First-strand synthesis kit (ThermoFisher Scientific, catalog number 18091050). 1 μl of prepared cDNA (equal to 50 ng of template RNA) was used in a 20 μl PCR reaction set up with Taq DNA polymerase. Annealing temperatures and cycle numbers were optimized using gradient PCR to the following conditions: primers for *FIN* and *FIN2* (table 1) were annealed at 52°C and 58°C, respectively, with 30 cycles for each PCR; for *FIN4,* primer (table 1) annealing temperature was optimized at 65°C for 25 PCR cycles. *Ubiquitin* was used as cDNA loading control using an equal quantity of template with ubiquitin primers (table 1) annealing at 53°C for 30 PCR cycles.

### FIN4 mRNA sequence, cloning and plant transformation

Full-length *FIN4* mRNA was amplified using forward and reverse primers designed based on the maximum extent of available mRNA sequence reads mapping to the *FIN4* region (table 1; Kudo et al., 2017; Lara-Mondragón et al., 2022). The resulting 1291 bp product was cloned into PCR II TOPO vector using TOPO TA cloning kit (ThermoFisher Scientific, catalog number 461020) and sequenced with T7 and SP6 primers to confirm the sequence of *FIN4* mRNA. The *FIN4* sequence intron and exon boundaries were different from those predicted by NCBI and Solgenomics, but when translated, produced a standard HPAT sequence (Supplemental Fig. 2).

To demonstrate rescue of the *fin4* mutant phenotype, 3,403 bp upstream of the *FIN4* start codon was cloned into pDONR221 with attB1 and attB5r recombination sites while the *FIN4* cDNA without stop codon was cloned into pDONR221 with attB5 and attB2 recombination sites using BP recombination (Gateway BP Clonase II Enzyme, ThremoFisher). The resulting entry vectors were combined by LR cloning reaction with the pFAST-R07 expression vector with the fluorescent protein mNeonGreen replacing GFP as a N-terminal fusion (Shimada et al., 2010; Beuder et al., 2020). The resulting plasmid containing *FIN4::FIN4cDNA-mNG* was transformed into cotyledon leaf explants of *fin4* plantlets via Agrobacterium-mediated transformation by the University of Nebraska Plant Transformation Core Research Facility.

### Protoplast transfection for cellular localization assay

For protoplast transfection, a Golgi-mCherry marker (the first 49 aa of GmMan1, soybean α-1,2-mannosidase I) obtained from the Nebenfuehr lab (Nelson et al., 2007) was cloned into the pEZS-NL vector with restriction cloning (table 1). The full-length *FIN4* cDNA without a stop codon was also cloned into pEZS-NL vector along with N-terminal fusion with mCerulean (table 1). Arabidopsis protoplast transfection was done as described in (Yoo et al., 2007) with 10 μg each of pEZS-NL Golgi mCherry and FIN4-CFP vectors. The transfected protoplasts were imaged 10 hours after transfection by Leica SP8 laser scanning confocal microscope.

### Histological staining and in situ hybridization

Paraffin embedding, toluidine blue staining and immunostaining were done as described previously (Jaffri and MacAlister 2021). For Alexander staining, pollen was collected from healthy dehiscent flowers and fixed in Carnoy’s fixative (6 ethanol:3 chloroform:1 acetic acid) for 2 hours. Fixed pollen was applied to a glass slide with a few of drops of simplified Alexander’s stain and imaged after applying a coverslip (Peterson et al., 2010). Stained pollen grains, were imaged using a Leica DM5500 compound microscope.

*In situ* hybridization was performed as described (Javelle et al., 2011). Briefly, 2 mm, 5 mm, and 8 mm flower buds were fixed and paraffin-embedded in RNase-free conditions. Full-length *FIN4* mRNA in PCRII-TOPO vector was used to synthesize sense and anti-sense probes using a DIG-labeled probe synthesizing kit (Millipore-Sigma, catalog number 11175025910) with T7 RNA polymerase for sense probe and SP6 RNA polymerase for antisense probe synthesis with 1μg plasmid as template. The synthesized probe was hydrolyzed by incubating at 60°C with 2X carbonate buffer (80 mM NaHCO_3_, 120 mM Na_2_CO_3_) for 52 minutes to produce 100-150 bp probes. After spin-column purification, 1μg of the probe was resuspended in 50μl 50% formamide and diluted 1:5 before applying to slides for hybridization. The probe was hybridized with 8μm thick tissue sections at 55°C for 20 hours. After two washes in 2X SSC buffer, slides were treated with NTE buffer (0.5 M NaCl, 10 mM Tris-HCl pH7.5, 1 mM EDTA pH8.0) containing RNase A (20μg/ml) for RNA digestion, followed by treatments with blocking buffer (0.5% blocking reagent in TBS buffer), washing buffer (1% BSA, 0.3% triton in TBS buffer) and then probed with alkaline phosphatase-conjugated anti-DIG antibody (diluted 1:5 in washing buffer) at 4°C overnight. After four washes with washing buffer, alkaline phosphate substrate solution was applied to each slide and refreshed twice daily for five days until the purple-blue color had developed. Slides were imaged using Leica DM5500 compound microscope after mounting the slides with Permount (Fischer Scientific, catalog number SP15-500).

### In vitro pollen germination and growth assays

For pollen tube length measurements, pollen collected from mature, dehiscent flowers was germinated in 4 ml of liquid pollen germination medium (PGM, Jahnen et al., 1989) in 6 well plates, in the dark. Pollen tubes were fixed at the appropriate time points by diluting PGM 1:1 with PGM containing 4% paraformaldehyde. Pollen tubes were imaged using a dissecting microscope scope at 6X magnification and measured using ImageJ software. For each genotype, 300 pollen tubes were measured per time point.

For pollen germination analysis, 500 μl of PGM was added to a 1cm x 1cm vacuum grease moat on a lysine-coated microscope slide. Pollen was then sparingly sprinkled with the help of a paintbrush and covered with coverslip. Pollen was considered to have germinated is an aperture protrusion was visible and was longer than the width of the pollen grain. At least 200 pollen were scored for each genotype and time point.

For pollen tube growth rate analysis, pollen was applied directly to a lysine-coated slide with a vacuum grease moat. Pollen germination media was then carefully added drop-wise to avoid displacing the pollen from the slide surface, and a cover slip was pressed to the moat to remove air space. Fields of ~30 pollen grains were imaged at 20X magnification, at five minute intervals for two hours. For each individual pollen tube, growth rate (μm/minute) was calculated for each five-minute interval by dividing the change in length in μm by the elapsed time. The average growth rate for each pollen tube was calculated by averaging the growth rate across all 24 time points.

For the pollen hydration assay, in a vacuum grease moated slide, a drop of pollen germination media was covered with a 1×1cm piece of dialysis membrane (Sigma-Aldrich catalog number, D9652-100FT). Fresh, dry pollen was sprinkled with a paintbrush and immediately sealed with a coverslip to avoid evaporation of PGM. The pollen were imaged using a Leica DM5500 compound microscope immediately and then every five minutes for an hour. Pollen was considered hydrated when the diameter around the middle of the grain increased, and the pollen became roughly spherical.

### Electron microscopy

For scanning electron microscopy (SEM), dry, mature pollen grains were collected from healthy dehiscent flowers and sprinkled on adhesive SEM stubs with paintbrush bristles. Pollen was sputter-coated with gold particles in a vacuum at 200 mAmps for 90 seconds for tomato pollen and 120 seconds for Arabidopsis pollen for optimum coverage. Gold-coated pollen was imaged using an EMAL JEOL JSM-7800FLV field-emission scanning electron microscope. 400-800 pollen grains were imaged and scored as collapsed if at least one distinct visible concave indentation was observed in one of the three inter-aperture regions.

Pollen grains were fixed and treated for transmission electron microscopy (TEM), as previously described (Jaffri and MacAlister, 2021). For quantification of wall layers, transverse sections of ten pollen grains of each genotype were imaged with three images taken for each of the three inter-aperture regions. Three measurements of wall thickness for each wall layer (sexine, nexine (layers of exine) and intine) were taken for each image and averaged.

## Results

### The hydroxyproline O-arabinosyltransferase FIN4 is expressed in anthers and pollen

Due to the importance of *HPAT1* and *HPAT3* in male fertility in Arabidopsis (MacAlister et al., 2016; Beuder et al., 2020), we hypothesized that members of this gene family might have a similar role in tomato pollen fertility. The tomato genome encodes four predicted *HPATs*, namely *FIN*, *FIN2, FIN3,* and *FIN4*. *FIN* has been previously shown to regulate meristem size through arabinosylation of members of the CLE peptide family, but no function has been assigned to the remaining tomato family members (Xu et al., 2015). We found that *FIN3* (Solyc07g021170) and *FIN4* (Solyc12g044760) were highly similar at the DNA-sequence level, sharing 88.3% base pair identity over more than 8 kilobases of sequence and are likely the result of an evolutionarily recent gene duplication (Supplemental Fig. 1a). Despite this sequence similarity, multiple attempts to clone a complete *FIN3* transcript were unsuccessful, suggesting it may be a pseudogene, or expressed under unique conditions. However, we successfully cloned the full coding sequence of the *FIN4* transcript and confirmed that it encoded a likely functional HPAT protein including a signal peptide, transmembrane domain and a highly conserved GT95 catalytic domain (Fig. 1a, Supplemental Fig. 2). HPATs, like many other glycosyltransferases, are Golgi-localized enzymes (Ogawa-Ohnishi et al., 2013; Xu et al., 2015). Sequence analysis of the cloned *FIN4* mRNA identified Golgi-localization signals in the predicted protein. To confirm FIN4 localization, we co-transfected Arabidopsis protoplasts with plasmids carrying C-terminal fusion of *FIN4* with mCerulean (mCer) as well as a Golgi-localized mCherry marker (the cytoplasmic tail and transmembrane domain of soybean α-1,2-mannosidase I) (Nelson et al., 2007). The co-localization of the FIN4-mCer signal with the signal from Golgi-mCherry confirmed that FIN4 is a Golgi-localized protein, consistent with the function of a canonical HPAT (Supplemental Fig. 1b).

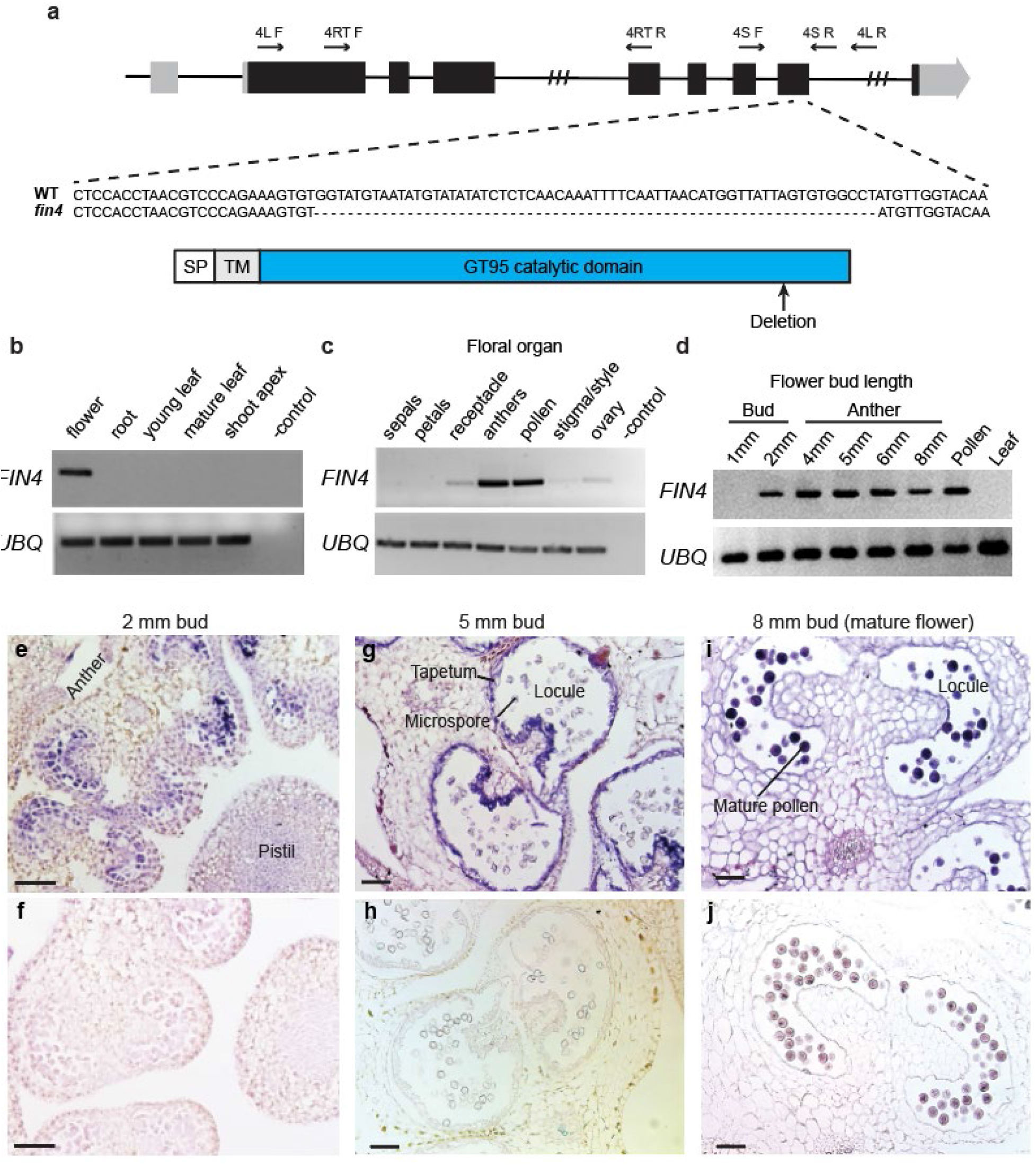
**a.** Diagram of the *FIN4* gene shows coding exons (black) and untranslated sequences (gray), CRISPR/Cas9 mediated deletion site, and the binding sites for primers used for genotyping and RT-PCR. Hash marks indicate truncated introns. Protein structure is shown below. SP is signal peptide, TM is transmembrane region. **b-d.** Semi-quantitative RT-PCR for *FIN4* by plant organ (**b**), floral organ (**c**), and developing flowers based on bud length (**d**). RNA was extracted from whole buds of the 1 mm and 2 mm bud stages and from developing anthers for 4, 5, 6, and 8 mm stages. Ubiquitin RT-PCR is used as a control. **e-j.** RNA *in situ* hybridization of developing anther sections probed with a *FIN4* antisense probe (e, g and i) and a *FIN4* sense probe (f, h and j). *FIN4* was expressed in the microspore mother cells of 2 mm bud anthers (e), in the uninucleate microspores and tapetal cells in 5 mm buds (g), and in the mature pollen of 8 mm buds (i). Scale bars = 25μm.

We next compared the expression patterns of the *FIN* genes using semi-quantitative RT-PCR of cDNA derived from different organs. We found that, while *FIN* and *FIN2* were broadly expressed across the tested organs (Xu et al., 2015; Supplemental Fig. 1c), *FIN4* transcript was limited to mature flowers (Fig. 1b), and within the flowers, it was most abundant in mature anthers and purified pollen (Fig. 1c). To determine the developmental timing of *FIN4* expression during floral development, we carried out semi-quantitative RT-PCR using floral tissue of multiple developmental stages. We have previously established a correlation between bud length and pollen developmental stage for the MicroTom tomato variety (Jaffri and MacAlister, 2021). We found *FIN4* was not expressed in floral buds preceding the specification of the microspore mother cells (1 mm buds), but was expressed in 2 mm buds containing microspore mother cells. Expression continued in developing anthers through the tetrad pollen stage (4 mm bud anthers), the uninucleate microspore stage (5 mm bud anthers), the bicellular pollen stage (6 mm bud anthers) and the mature pollen stage (8 mm bud anthers) (Fig. 1d). To establish tissue specificity in this expression timeline, we used RNA *in situ* hybridization to detect *FIN4* transcripts in anther sections (Fig. 1e-j). In 2 mm buds, we observed staining in pollen mother cells (Fig. 1e). In 5 mm buds, we detected *FIN4* transcript in the uninucleate microspores as well as the sporophytic tapetal cells lining the locule cavity (Fig. 1g). In mature (8 mm bud) anthers in which the tapetum had degenerated, we detected *FIN4* transcript in the mature pollen grains (Fig. 1i). These results show that within the anther, *FIN4* mRNA transcript was enriched in the pollen precursors and tapetal cells. Given *FIN4’*s expression profile, we hypothesized that it had a role in male fertility in tomato, possibly similar to the function of *HPAT1* and *HPAT3* in Arabidopsis.

### Loss of FIN4 impairs male fertility but not pollen tube elongation

To investigate *FIN4’s* contribution to male fertility, we obtained seeds for a previously generated CRISPR/Cas9-induced *FIN4* deletion mutant (Brooks et al., 2014). The mutant *fin4* allele is a 68 bp deletion in the 7th exon, triggering a frameshift in the highly conserved catalytic domain, ultimately disrupting the last 30 amino acids of the protein (Fig. 1a; Supplemental Fig. 2). The mutant was out-crossed to MicroTom to remove the CRISPR/Cas9 transgene followed by two additional generations of out-crossing to standardize the genetic background. The resulting BC3 generation *fin4/+* plants were allowed to self-fertilize to assess transmission of the mutant allele. Despite robust seed set, of the 160 genotyped progeny, only one (0.6%) was homozygous for *fin4,* a significant deviation from the expected 25% for a Mendelian segregation pattern (N=160, χ^2^ P-value=1.0762×10^-12^). The homozygous *fin4* plant was morphologically normal and was able to produce viable seed, suggesting a transmission defect rather than a developmental defect in the homozygous *fin4* plants. To determine if the mutant allele transmission was disrupted through the male, the female or both, we performed a series of test crosses between *fin4/+* and WT. In the absence of a transmission defect, the progeny of such a cross is equally likely to inherit the WT or *fin4* allele from the *fin4/+* parent, resulting in an expected 1:1 ratio of WT and *fin4/+* progeny. When *fin4/+* used as the female parent, we observed the expected transmission pattern (15 *fin4/+* and 17 WT; χ^2^ P-value=0.7237), indicating that female transmission was not altered. However, when *fin4/+* was used as the pollen parent, only 6.25% of the resulting progeny inherited the mutant allele, a statistically significant deviation from expectations (2 *fin4/+* and 30 WT; χ^2^ P-value=7.43×10^-7^), consistent with a pollen-specific fertility defect.

Arabidopsis *hpat1 hpat3* double mutants also display reduced male transmission due to impaired pollen tube elongation and high frequencies of pollen tube rupture, leading to shorter pollen tubes (MacAlister et al., 2016). To determine if *fin4* pollen behaves similarly, we compared lengths of *in vitro* grown pollen tubes between WT and *fin4.* After three hours of *in vitro* growth, the *fin4* pollen tubes had a minor, but statistically significant reduction of pollen tube length compared to the WT pollen tubes (Fig. 2b). However, unlike *hpat1 hpat3* pollen tubes which frequently display morphological defects, the morphology of the *fin4* pollen tubes was normal (Fig. 2a-b). Furthermore, when we measured the growth rate of individual pollen tubes we found no significant different between WT and *fin4* pollen (Fig. 2c). Therefore, unlike the Arabidopsis *hpat1 hapt3* phenotype which traces to defects in pollen tube elongation, the tomato *fin4* pollen tubes have a similar growth competence compared to those of WT.

**Figure 2.**
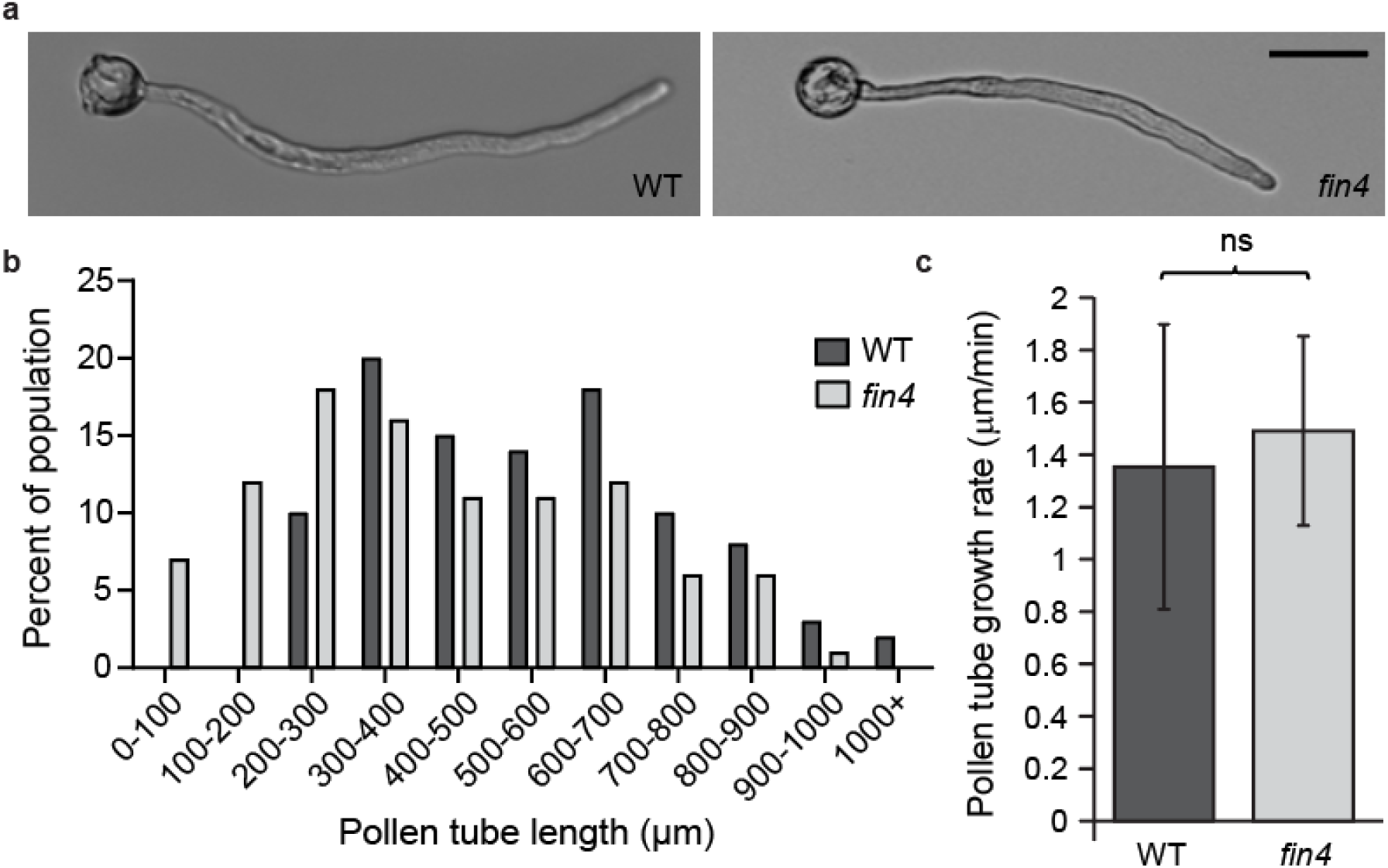
Loss of *FIN4* does not prevent pollen tube elongation. **a.** Representative *in vitro* grown WT (left) and *fin4* (right) pollen tubes. Scale bar = 50μm. **b.** Pollen tube length-frequency distribution after three hours of *in vitro* growth, (mean WT 534.4±11.36μm, mean *fin4* 474.1±14.85μm, mean±SD, N=300 pollen tubes/genotype; T-test P-value=0.0013). **c.** Average growth rate of WT and *fin4* pollen tubes in μm/minute. Ns indicates no statistically significant difference between the two genotypes.

### *Though viable,* fin4 *pollen frequently fail to hydrate and germinate*

Given that the reduced male transmission of *fin4* was not adequately explained by *in vitro* pollen tube growth defects, we next examined other potential explanations for the low fertility of *fin4* pollen. Compromised pollen transmission may be caused by several defects, including the release of aborted or otherwise non-viable pollen from the anther. To assess pollen viability, we used Alexander stain, a differential pollen viability stain on released pollen and counted the proportion of apparently non-viable pollen (Peterson et al., 2010). We found high levels of viability in both the WT and *fin4* pollen with no statistically significant difference between genotypes (Supplemental Fig. 3) (N>1100 pollen/genotype; t-test P-value=0.1950). Therefore, the pollen produced by *fin4* plants have cytoplasmic content that is grossly similar to that of the WT plants.

Interestingly, despite the apparent high viability of *fin4* pollen grains, we noted frequent failure to germinate and initiate pollen tube growth during *in vitro* growth assays (Fig. 3a-b). When we measured pollen germination over time, we found a statistically significant reduction in *fin4* pollen germination at each time point measured (Fig. 3c). While WT pollen reached ~90% germination within two hours in pollen growth media, *fin4* pollen reach a maximum of ~23% germination at this time (N>200 pollen/genotype; χ^2^ P-value= 1.138×10^-70^). Given that *FIN4* was expressed in both the developing pollen and in the sporophytic tapetal cells (Fig. 1g), we also measured the germination ability of pollen from *fin4/+* plants to determine if the germination defect was specific to the genotype of the pollen itself or a consequence of development in the *fin4* sporophytic context. Half of the pollen produced by a *fin4/+* plant will be genetically wild-type and the other half will carry the *fin4* mutant allele though all of the pollen will share the same heterozygous sporophyte genotype. Consistent with the pollen genotype determining germination capacity, the germination of the *fin4/+* pollen was intermediate between that of *fin4* and WT (Fig. 3c).

**Figure 3.**
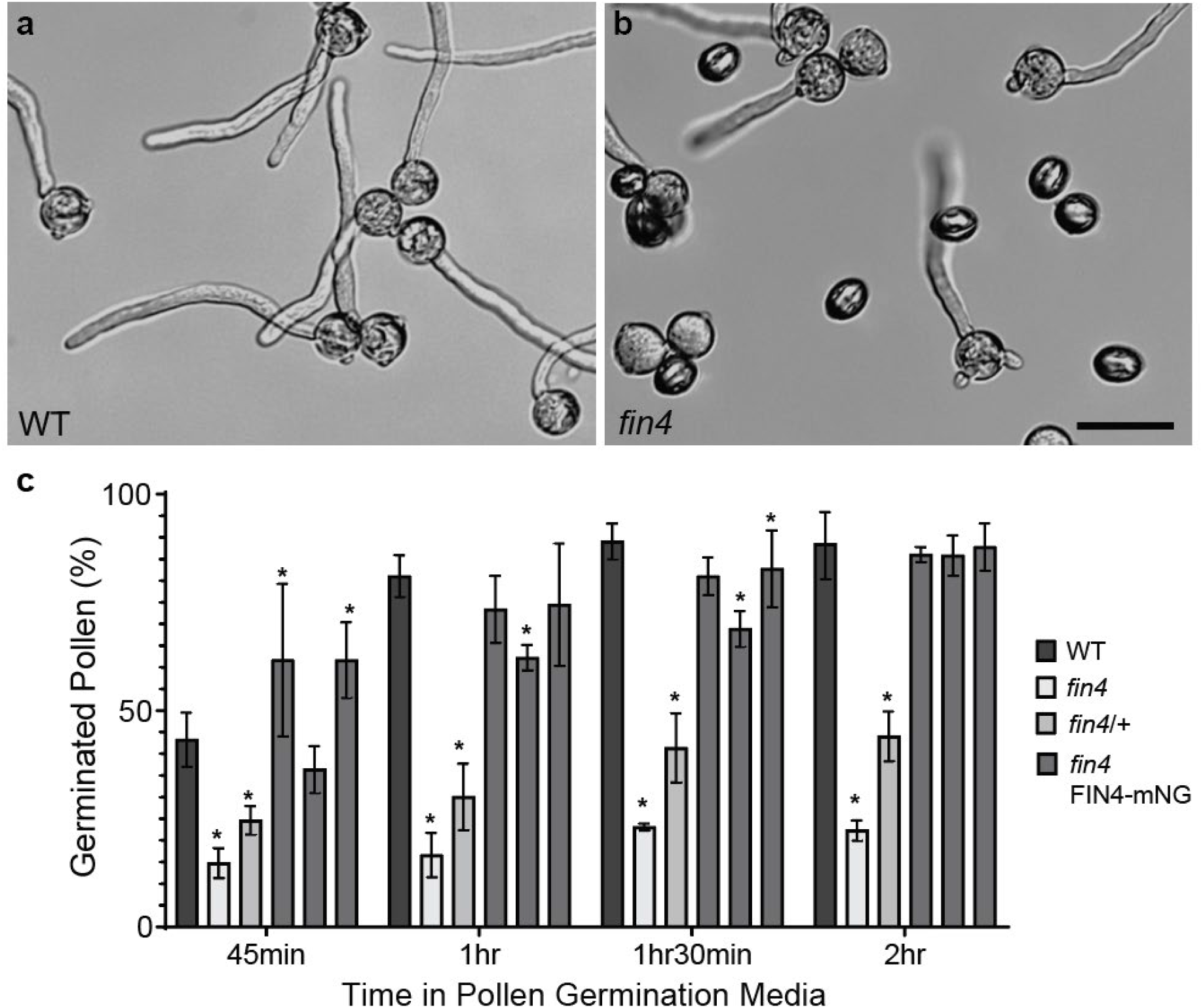
*fin4* pollen has reduced germination. Two hour *in vitro* grown pollen of WT (**a**) and *fin4* (**b**). **c.** Quantification of pollen germination over time (mean ± standard deviation for three independent experiments). Asterisks mark statistically significant differences compared to WT (Bonferroni corrected χ^2^ P-value <0.05).

To confirm that the low germination of *fin4* mutant pollen was caused by loss of FIN4 activity, we transformed *fin4* plants with a transgenic rescue construct containing 3.7 kb of the native *FIN4* promoter, the *FIN4* cDNA sequence, and a C-terminally fused mNeonGreen fluorescent protein (FIN4-mNG). In three independent transgenic lines, we found pollen germination was restored to near wild-type levels with no statistically significant difference between the transgenic lines and WT after two hours in germination media (Fig. 3c). While this demonstrates that the transgenic protein is functional and that the *fin4* mutant phenotype is due to a loss of FIN4 protein function, unfortunately, auto-fluorescence within the anther prevented satisfactory imaging of the FIN4-mNG fusion protein *in vivo.*

Pollen hydration is a necessary precursor to pollen germination and is associated with an increase in volume and a change in pollen shape from elongated to spherical. During our *in vitro* pollen germination assays, we frequently observed *fin4* pollen failed to change shape, indicating a failure to hydrate (Fig. 4a). To more closely mimic the hydration environment of the tomato stigma surface, we scored pollen hydration on a semi-permeable membrane (Wolters-Arts et al., 2002). Most wild-type pollen (88.9%) successfully hydrated within one hour with many (40.8%) hydrating within 10 minutes. However, only 13.8% of *fin4* pollen achieved hydration within one hour, a significant reduction compared to WT (N>100/genotype; χ^2^ P-value=1.94×10^-29^) (Fig. 4). Therefore, though viable, *fin4* pollen have significantly reduced ability to successfully hydrate and initiate pollen tube growth. However, once a pollen tube has been initiated, it is able to elongate similarly to WT pollen tubes.

**Figure 4.**
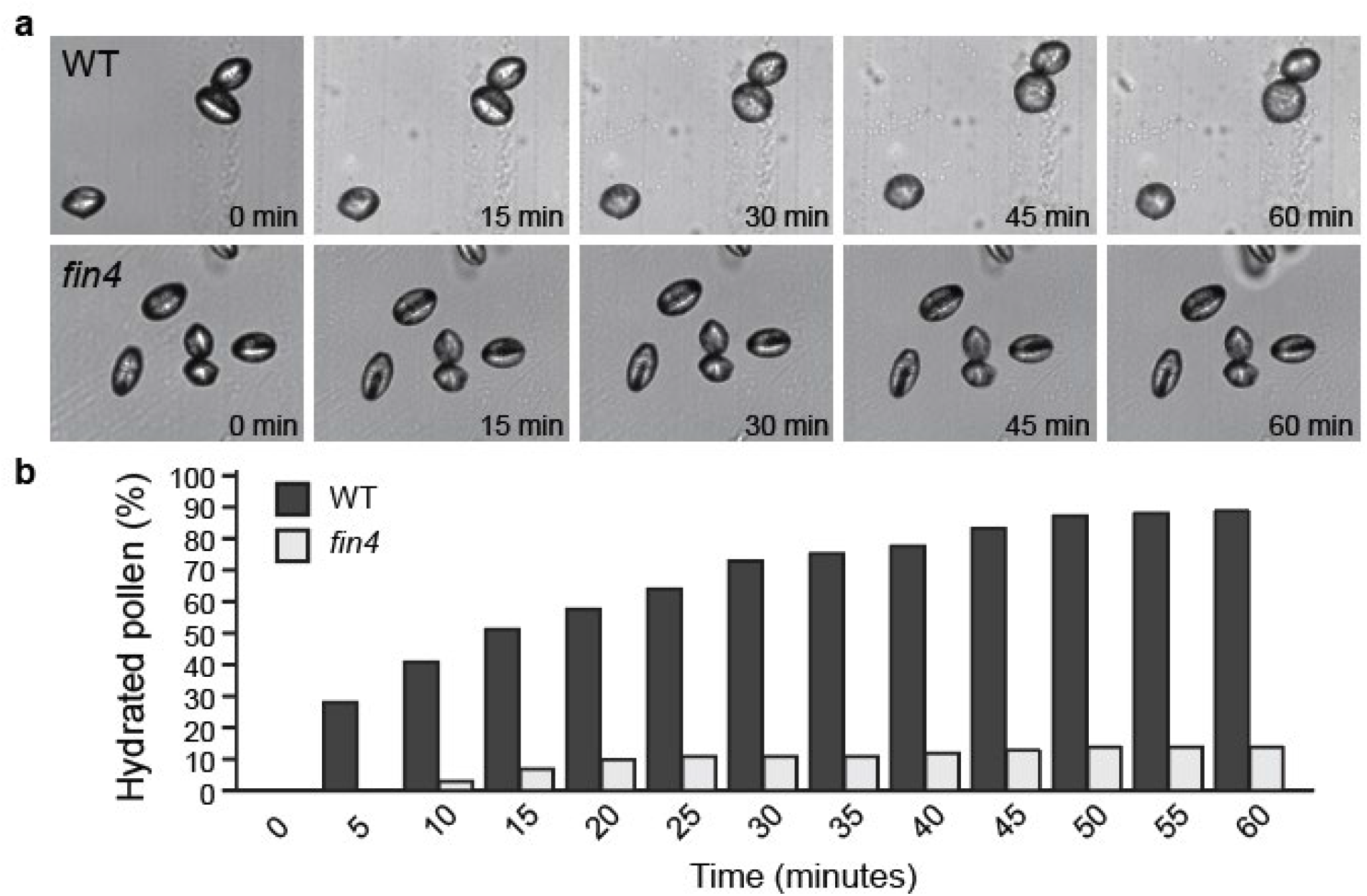
*fin4* pollen largely fail to hydrate. **a.** Time-lapse of WT pollen (top) and *fin4* pollen (bottom) on semi-permeable membrane. Shape change from elongate to spherical indicates successful hydration. **b.** Quantification of pollen hydration of WT and *fin4* pollen at five-minute intervals (N>100/genotype; χ^2^ P-value=1.94×10^-29^)

### fin4 *pollen grains have compromised structural integrity*

Given the compromised hydration ability of *fin4* pollen, we hypothesized that a structural defect in the pollen wall might prevent the passage of water into the pollen grain. To observe the external structure of mature pollen grain, we used scanning electron microscopy to image the pollen. While we found no visible difference in the appearance of the exine (Fig. 5a-b), we frequently observed collapsed *fin4* pollen following SEM sample preparation (Fig. 5c-g; 61.0% of *fin4* vs. 9.9% of WT pollen; N>400-800 pollen/genotype; χ^2^ P-value = 1.1032×10^-99^). Since the *fin4* pollen shape was normal under standard light microscopy conditions, we interpret the *fin4* pollen collapse as a response to the physical stress of SEM sample preparation which involves sputter-coating dry pollen with gold particles under vacuum. This apparent difference in the ability to resist the physical stress of gold coating suggested differences in the mechanical properties of the *fin4* and WT pollen grains. As was the case with the germination phenotype, the phenotype of pollen from *fin4/+* plants was intermediate between that of *fin4* and WT with 32.8% of the pollen being collapsed. The SEM collapse phenotype was also reduced to the level of the WT in the *fin4* FIN4-mNG transgenic lines (Fig. 5g).

**Figure 5.**
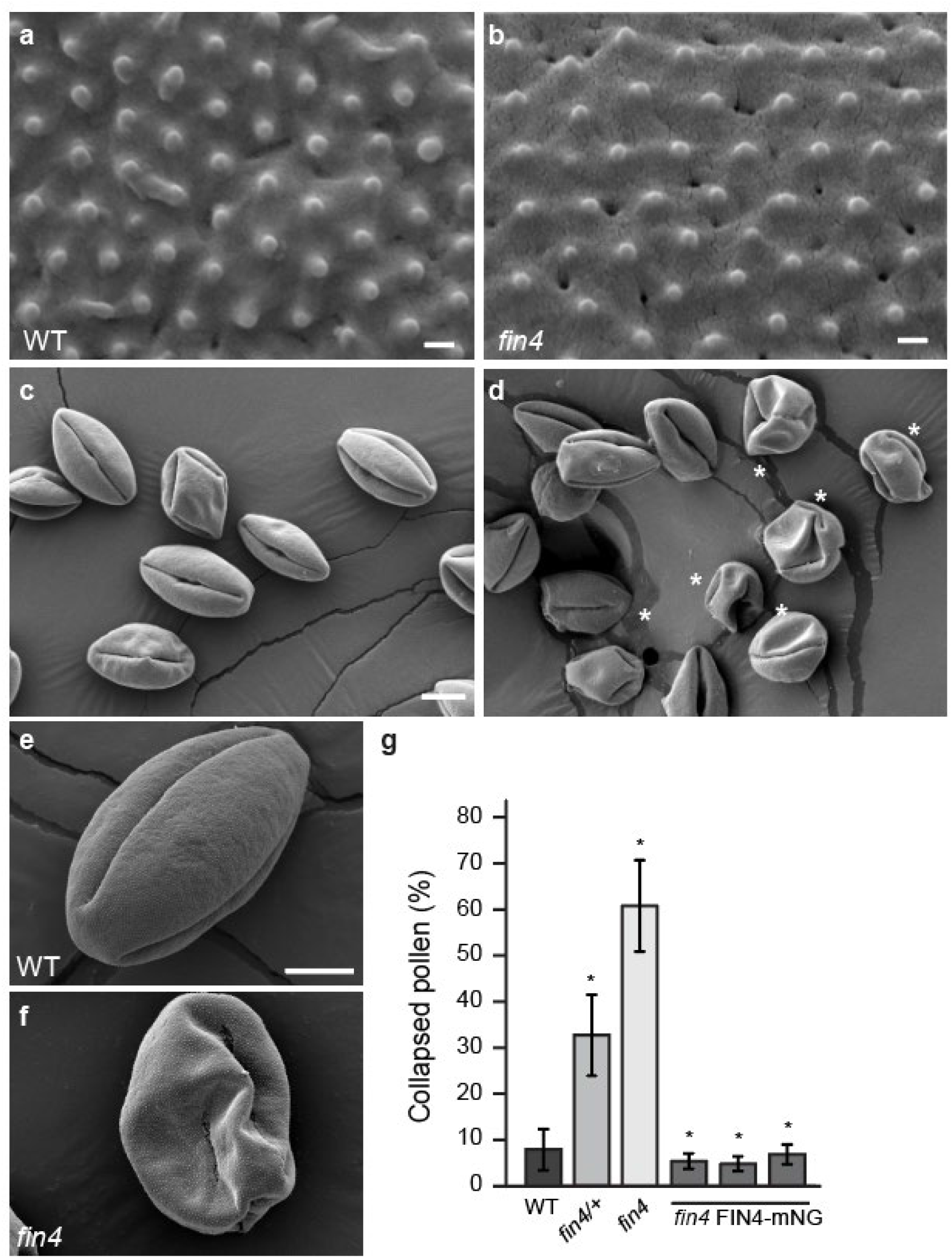
*fin4* pollen are prone to collapse. Scanning electron micrographs (SEM) of WT (**a**) and fin4 (**b**) pollen surface showing exine patterning. Scale bars in a and b = 100 nm. **c-f.** SEM of mature, released pollen grains following gold coating. While WT pollen generally maintain their shape following SEM sample preparation (**c** and **e**), *fin4* pollen are prone to collapse (**d** and **f**), asterisks mark collapsed pollen grains, scale bars in c-f = 600 nm. **g.** Quantification of the frequency of pollen grain collapse following SEM sample preparation.

Given the novelty of this phenotype, we also subjected Arabidopsis *hpat1 hpat3* and WT Columbia-0 pollen to SEM sample preparation. We observed no visible difference between the Columbia-0 and the *hpat1 hpat3* mutant, with no collapsed pollen observed for either genotype (Supplemental Fig. 4), confirming that *FIN4’s* function in tomato pollen fertility differs from that of *HPAT1* and *HPAT3* in Arabidopsis.

### *Intine formation is disrupted in* fin4 *pollen*

To better understand the structural differences between *fin4* and WT pollen, we used transmission electron microscopy (TEM) to observe cross sections of mature pollen grains, focusing on pollen wall structure. Initial comparison between WT and *fin4* pollen showed that, consistent with the viability staining results (Supplemental Fig. 3), the cytoplasmic content was not noticeably different between WT and *fin4* pollen (Fig. 6a-b). However a comparison of the pollen grain wall indicated differences between the two genotypes (Fig. 6c-d). While the exine layers (both the continuous nexine and the sculpted sexine) were similar in thickness and appearance between wild-type and *fin4* pollen, the intine thickness was significantly reduced in *fin4* (Fig. 6e-g), with some regions apparently lacking intine completely (Fig. 6h). These results indicate that while the amount of exine deposition did not differ between WT and *fin4* pollen, the amount of intine was significantly reduced in mature *fin4* pollen.

**Figure 6.**
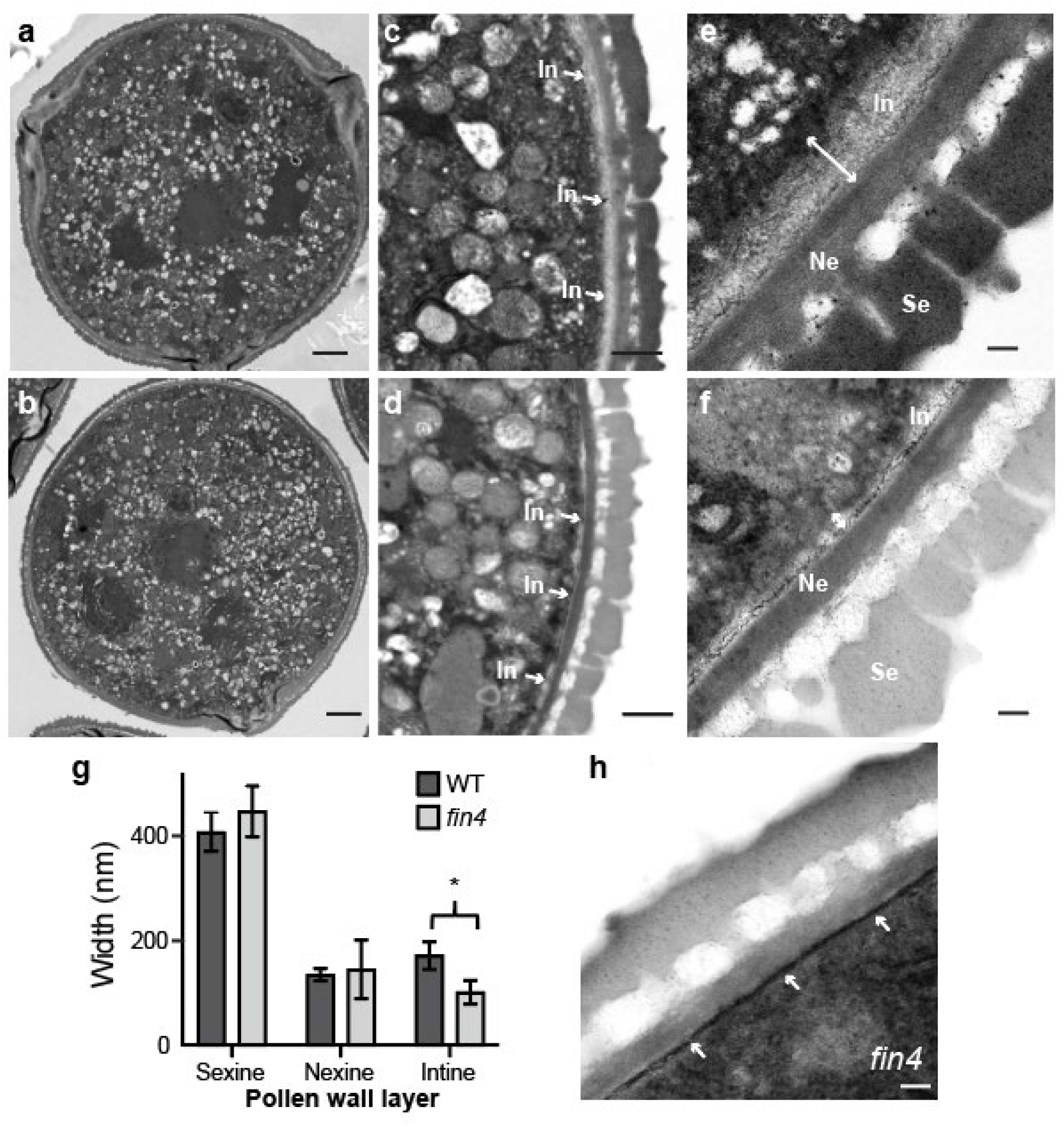
The intine is reduced in *fin4* pollen. TEM of mature pollen grains showing the cross-sectional structure of the released pollen wall of WT (**a**, **c** and **e**) and *fin4* (**b**, **d**, **f** and **h**). Sections showing complete pollen grains (**a** and **b,** scale bars = 2μm) and magnified views of inter-aperture region of pollen grain wall (c-f, scale bars = 600 nm in c and d and 100 nm in e, f and h) with the lighter intine layer (In) indicated with arrows and the layers of the exine (sexine, Se and nexine, Ne) layers marked in **e** and **f.** Quantification of pollen wall thicknesses in WT and *fin4* pollen (**g**, mean ±SD, ‘*’ marks statistically significant difference, T-test p-value <0.05). Some areas in the *fin4* pollen show apparently complete intine loss as marked by the arrows (**h**).

To better understand the developmental origin of the reduced *fin4* intine, we examined pollen at earlier developmental stages. We have previously established that in 6 mm MicroTom flower buds, pollen is at the bicellular stage of development and intine deposition has begun, particularly at the aperture sites and in 8 mm buds the pollen is nearly mature and the intine is well established throughout the pollen (Jaffri and MacAlister, 2021). To understand the nature of the *fin4* intine defect, we used calcofluor white, a fluorescent dye with a high affinity for cellulose (β-1,4-glucan), often used to visualize the pollen intine layer to label sections from 6 and 8 mm flower buds (Fang et al., 2008; Herburger & Holzinger, 2016; Jaffri & Macalister, 2021; W. L. Li et al., 2017; Renzaglia et al., 2020; Takebe et al., 2020). As we had previously found, WT pollen in the 6 mm buds had begun intine deposition, visible as calcofluor white fluorescence, particularly accumulating at the future aperture sites (Fig. 7a). In the mature pollen of 8 mm WT buds, the calcofluor white staining had formed a continuous ring around the pollen grain with noticeable accumulation at the aperture sites (Fig. 7c). In *fin4* mutant pollen however, we noted a major deviation from this pattern of accumulation. While the surrounding anther tissue was well stained with calcofluor white, the developing pollen grains were largely missing detectable fluorescence at both the 6 and 8 mm bud stage (Fig. 7b and d), suggesting that intine development, specifically cellulose deposition was impaired in the *fin4* pollen leading to a reduction in this specific wall layer in the mature pollen grains.

**Figure 7.**
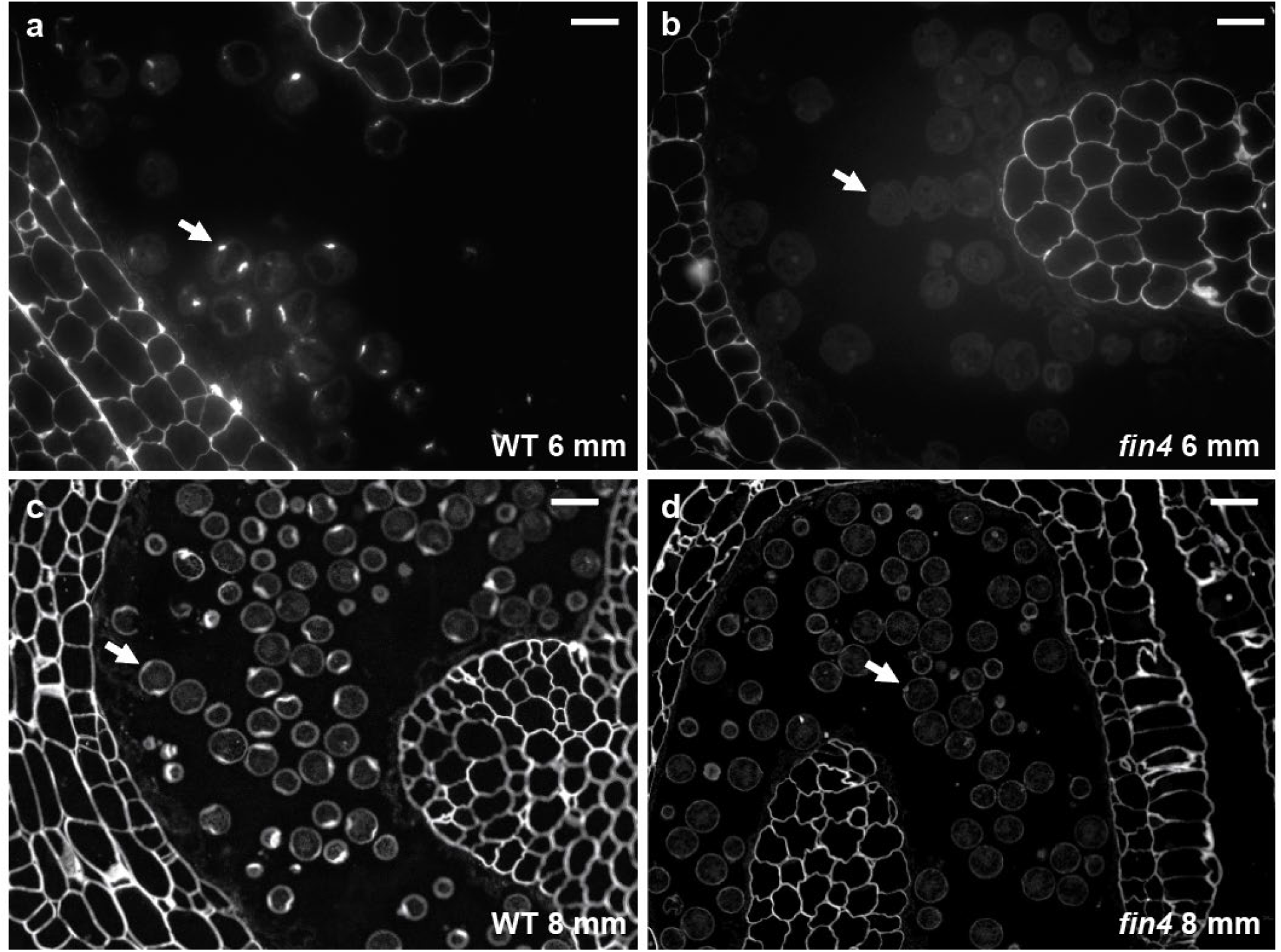
Cellulose deposition is reduced in developing *fin4* pollen intine. Cross section of anthers from 6 mm (bicellular pollen stage, **a** and **b**) or 8 mm buds (mature pollen stage, **c** and **d**) stained with calcofluor white to detect cellulose. Cellulose returns to the pollen wall first at the incipient aperture regions in WT bicellular pollen (**a**) but is not detectable in *fin4* at this stage (**b**). As pollen development continues, the remainder of the intine is formed and cellulose is detectable around the periphery of the WT pollen (**c**). Cellulose detection remains weaker in *fin4* pollen (**d**). Arrows mark individual pollen grains. Scale bars = 100 μm.

Several lines of evidence suggest that pectin dynamics play an important role in pollen intine development (Cankar et al., 2014; Chebli et al., 2012; Jiang et al., 2013; Phan et al., 2011; Suárez-Cervera et al., 2002). Given the hypothesized function of the EXT proteins as a scaffold for pectin deposition (Cannon et al., 2008) and the role of HPATs in EXT glycosylation (Ogawa-Ohnishi et al., 2013), we next sought to determine if pectin distribution was also disrupted during intine deposition in *fin4* pollen. The monoclonal antibodies LM20 and LM19 bind to HG with a low (dmeHG) and high (meHG) degree of esterification, respectively (Christiaens et al., 2011; Verhertbruggen et al., 2009). In 6 mm WT buds we observed a diffuse LM20 (meHG) signal around the periphery of the bicellular pollen (Fig. 8a) while in *fin4* pollen, the signal was much more robust (Fig. 8b). The binding of LM19 (dmeHG) followed a similar pattern wherein WT pollen signal was faint with some additional accumulation at the developing aperture sites (Fig. 8c). In *fin4* pollen, we again found robust LM19 binding around the pollen periphery (Fig. 8d). Thus, while cellulose production was apparently reduced, HG deposition appeared to have increased during intine formation.

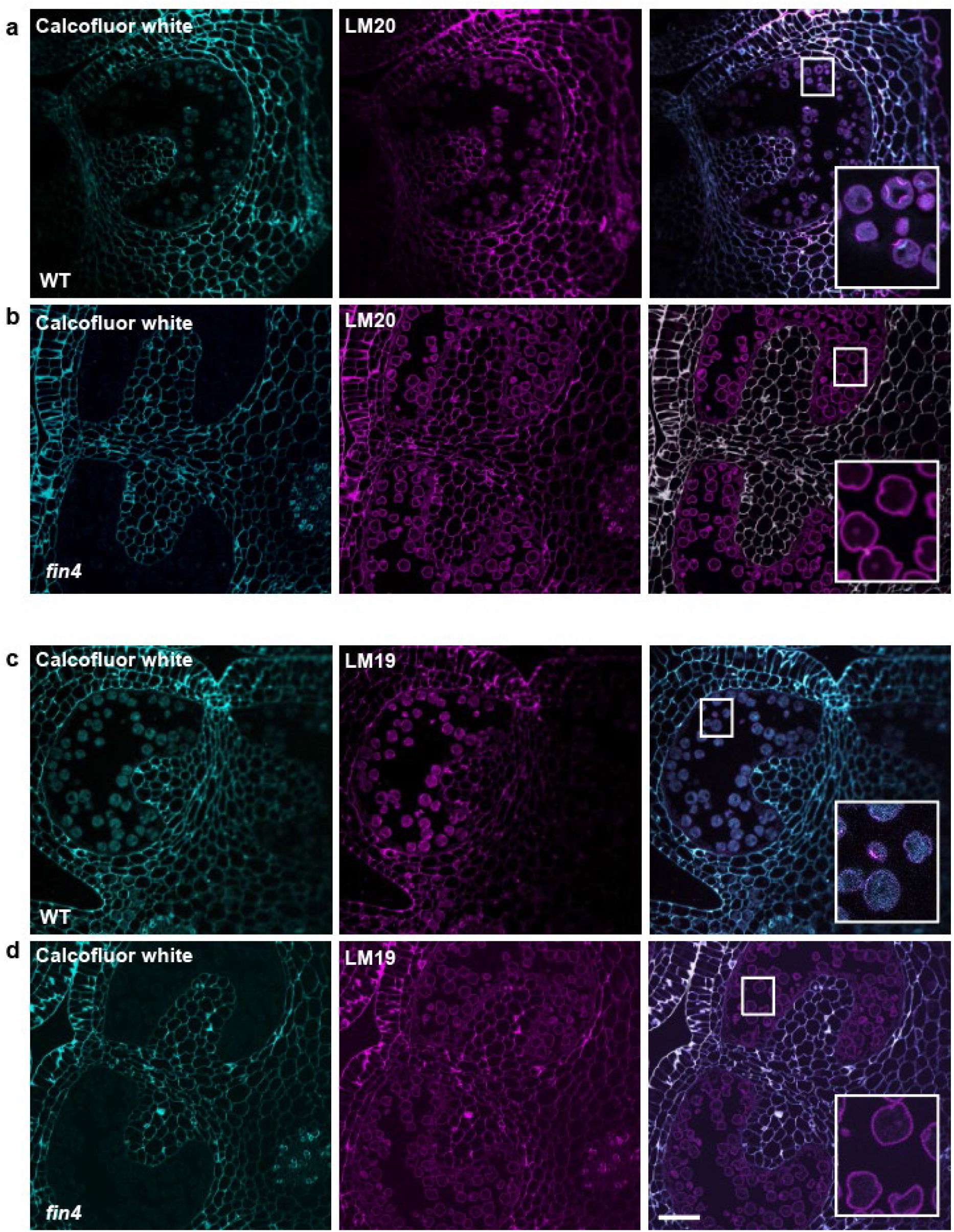
Fluorescent micrographs of anther cross-sections of 6 mm bud of WT (**a** and **c**) and *fin4* (**b** and **d**) stained with Calcofluor white (teal, left panels) and LM20 or LM19 primary antibody detected with FITC-conjugated secondary antibody (magenta, center panels) and merged channels (right panels) with inset showing higher magnification of the indicated region. Scale bars = 50μm.

## Discussion

Pollen has a highly complex and unique cell wall organization with a correspondingly complex developmental pathway. The formation of the inner intine layer is particularly poorly understood. Here we show that tomato mutants of the hydroxyproline *O*-arabinosyltransferase, *fin4* have a severe and specific defect in intine formation during the late stages of pollen development. Aside from significantly reduced intine deposition, *fin4* pollen appeared cytologically normal and viable (Fig. 6), but frequently failed to hydrate and germinate (Fig. 3 and 4), leading to significantly reduced fertility as measured by mutant allele transmission.

*FIN4* is likely to encode a functional HPAT enzyme due to its complete and highly conserved GT95 catalytic domain and Golgi localization (Supplemental Fig. 1B and 2). Therefore, the *fin4* phenotype is likely due to loss of hydroxyproline *O*-arabinosylation of one or more key target proteins that support intine formation. HPATs are known to modify cell wall-associated proteins, specifically EXTs (Ogawa-Ohnishi et al., 2013). The *EXT* family is generally large and subject to functional redundancy and complex transcriptional feedback, making the study *EXT* function complex (Choudhary et al., 2015). The tomato genome encodes 83 *EXT* members, including several canonical *EXTs* as well as multiple classes of genes encoding *EXT*-like regions combined with other domains (e.g. Leucine-rich repeat EXT-like proteins (LRXs), Proline-rich extensin-like receptor kinases (PERKs) and Class I Formin Homology proteins (FHs); Ding et al., 2020). Several *LRX* and *FH* genes have been implicated in pollen tube growth in Arabidopsis (Borassi et al., 2016; Sede et al., 2018; Lara-Mondragón et al., 2022; Mecchia et al., 2017; Wang et al., 2018). A role in intine formation has also been briefly described for some Arabidopsis LRX members. In TEM images, *lrx8/9* and *lrx8/9/10/11* mutant intine contains a central electron dense band rather than the continuous, light intine observed in WT pollen grains (Fabrice et al., 2018). Furthermore, immunolocalization of a maize LRX protein, Pex1, demonstrated intine localization in mature pollen grains, suggesting a potential general function in intine structure, though the molecular basis of this function is unknown (Rubinstein et al., 1995). Following their glycosylation, EXTs are secreted into the cell wall space where they may form a covalently cross-linked network which is hypothesized to serve as a scaffold for the assembly of cell wall polymers, specifically pectins (Cannon et al., 2008; Nuñez et al., 2009). Deglycosylated EXTs fail to properly crosslink *in vitro,* suggesting that loss of glycosylation caused by absence of HPAT activity could inhibit proper scaffold assembly (Chen et al., 2015). Given the hypothesized relationship between EXTs and pectin, we examined the abundance and distribution of HG epitopes during intine development. While we found reduced calcofluor white binding indicating a reduction in cellulose content in the *fin4* pollen wall (Fig. 7), we observed in increase in both LM19 and LM20 binding in the mutant pollen, suggesting increased levels of dmeHG and meHG, respectively (Fig. 8). Alternatively, the increase in LM19 and LM20 binding may be explained by an “unmasking” of epitopes of these antibodies due to other cell wall changes (e.g. failure to associate with EXTs) allowing more efficient antibody binding without a change in absolute polymer abundance. However, changes in pectin are consistent with the *fin4* hydration defect (Fig. 4). Pectin de-methylesterification is required for normal pollen hydration as demonstrated by Arabidopsis *pme48* mutants which display higher degrees of HG methylesterification and significantly delayed pollen hydration and germination (Leroux et al., 2015). Similarly, chemically de-esterified pollen particles from sunflower show pH dependent swelling responses that are regulated by calcium abundance (Fan et al., 2020). Interestingly, *Brassica campestris pme37c* mutants also have altered intine formation, however, unlike *pme48* mutants display an apparent over-hydration phenotype of some pollen grains (i.e. a greater in increase in pollen diameter than that of WT following hydration; Xiong et al., 2019). Pectin status, therefore, plays a key functional role in pollen hydration and *fin4’s* pollen hydration defect may be due to disrupted phase transitions within the intine caused by altered intine composition (Haas et al., 2021). Determining how *FIN4* influences pectin deposition and the specific glycosylation targets required for intine formation, whether EXTs, EXT-like proteins, or other types of target proteins, will require further research.

Like *fin4* mutants, Arabidopsis *hpat1 hpat3* double mutants also have reduced pollen fertility (MacAlister et al., 2016). However, the *hpat1 hpat3* fertility defect traces to compromised pollen tube growth (slower pollen tube elongation and high incidence of pollen tube rupture) rather than defective pollen hydration and germination (Beuder et al., 2020). The approximately 20% of *fin4* pollen that successfully germinated during *in vitro* assays (Fig. 3c) formed morphologically normal pollen tubes that elongated at the same rate as those of the WT (Fig. 2a and c). Therefore, while the *Arabidopsis HPAT*s are important for organizing the cell wall of the pollen tube, *FIN4* functions earlier during pollen intine development and is not required for pollen tube growth after germination. Like the *fin4* intine, the composition of the pollen tube walls in *hpat1 hpat3* is disrupted. In pollen tubes, the distribution of cell wall polymers is highly polarized with “reinforcing” polymers like callose (β-1,3-glucan) and dmeHG excluded from the expanding, meHG-rich pollen tube tip (Chebli et al., 2012; Cascallares et al., 2020). In *hpat1 hpat3* pollen tubes this polarized distribution is disrupted with callose and dmeHG invading the tip region, consistent with the reduced growth rates of the mutant tubes (Beuder et al., 2020). A genetic suppressor screen for secondary mutations capable of increasing *hpat1 hpat3* pollen fertility identified secretory pathway genes belonging to the exocyst complex and the Sec1/Munc18 gene *SEC1A* (Beuder et al., 2020; Beuder et al., 2022). Reduced rates of HPAT-targeted glycoprotein secretion in the suppressed pollen tubes led to a partial restoration of the cell wall organization and improved pollen tube growth, suggesting that HPAT-dependent glycosylation is important for maintaining cell wall structure.

## Acknowledgements

We would like to thank Prof. Tom Clemente and the University of Nebraska Plant Transformation Core Research Facility for transformations, Erik Nielsen for providing the Golgi localization reporter, Gregg Sobocinski for imaging advice and Zach Lippman for providing the *fin4* seeds. This work was supported by the National Science Foundation under grant no. IOS-1755482.

## Supplemental Material

**Supplemental Figure 1.**
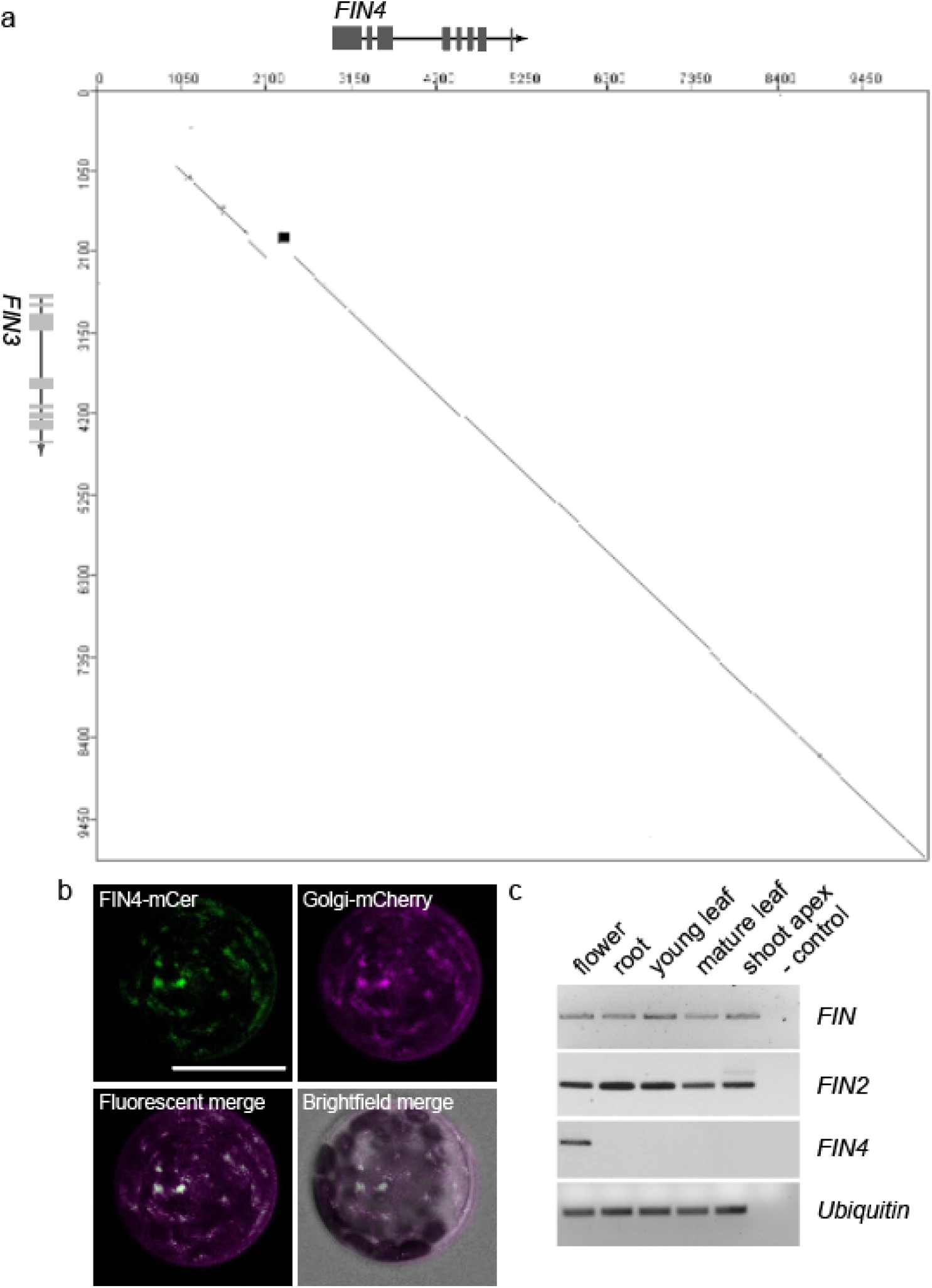
*FIN4* genomic context, protein localization and gene expression pattern. **a.** Dot blot graphic summary of *FIN3* and *FIN4* genomic region alignment, showing the region of the genome duplication. **b.** Fluorescent micrograph of Arabidopsis mesophyll protoplasts cotransfected with plasmids carrying C-terminal FIN4-CFP (green) and a Golgi localized mCherry marker (magenta) as well as fluorescent merged and bright field merged images. Scale bar = 1 mm. **c.** Semi-quantitative RT-PCR for *FIN, FIN2* and *FIN4* by plant organ. Shoot apex includes vegetative meristem and leaf primordia. Ubiquitin RT-PCR is used as a loading control. While *FIN* and *FIN2* are broadly expressed, FIN4 expression was limited to mature flowers.

**Supplemental Figure 2.**
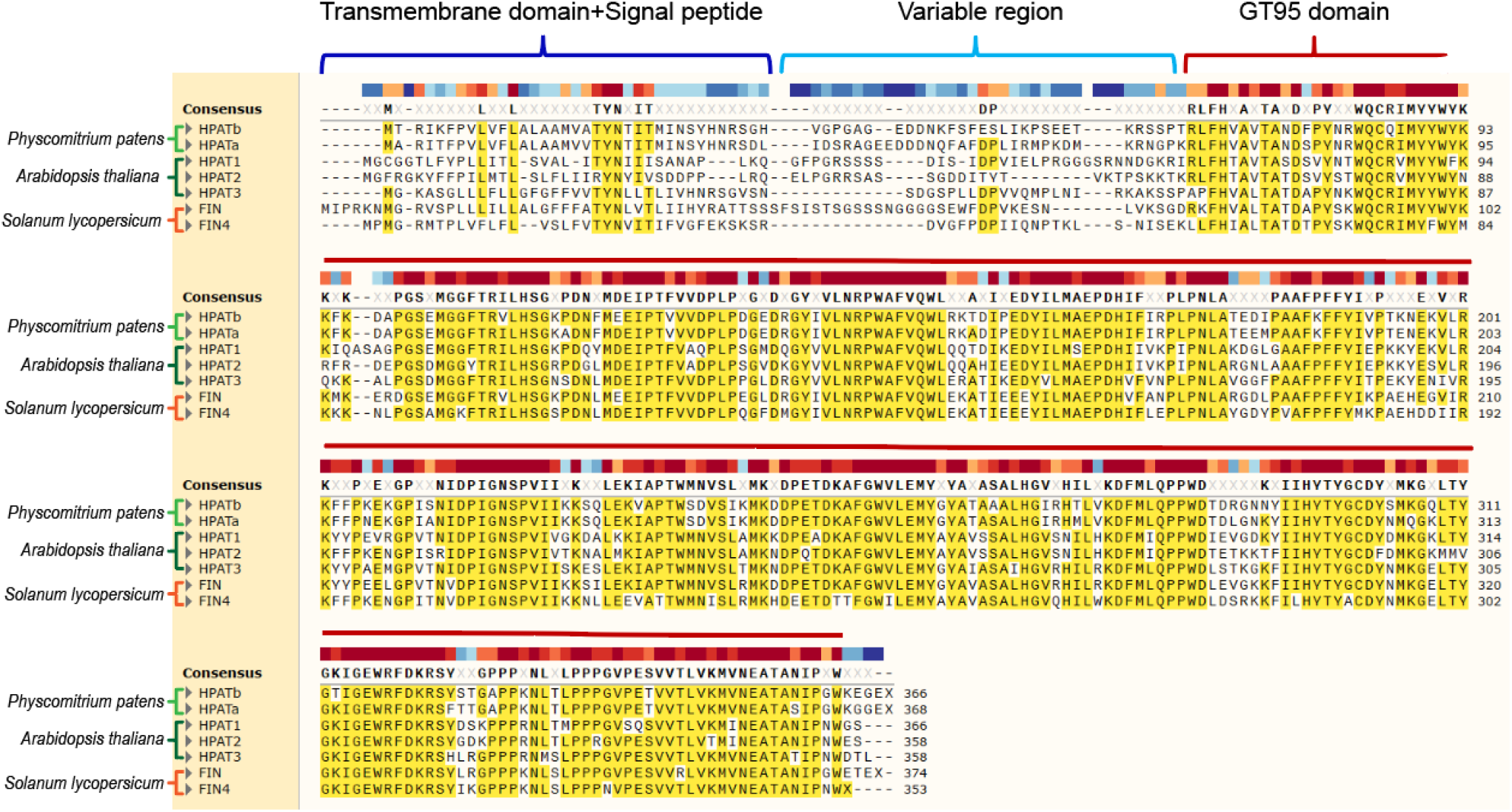
Protein sequence alignment of select HPATs. Graphic summary of protein sequence alignment of FIN and FIN4 with HPATs from Arabidopsis and the model moss *Physcomitrium patens* showing the consensus sequence above. Major protein regions are marked including the transmembrane domain and signal peptide, the variable (weakly conserved) region, and the highly conserved portion corresponding to the GT95 catalytic domain. The heat map of alignment shows blue for low conservation to dark red for most conserved.

**Supplemental Figure 3.**
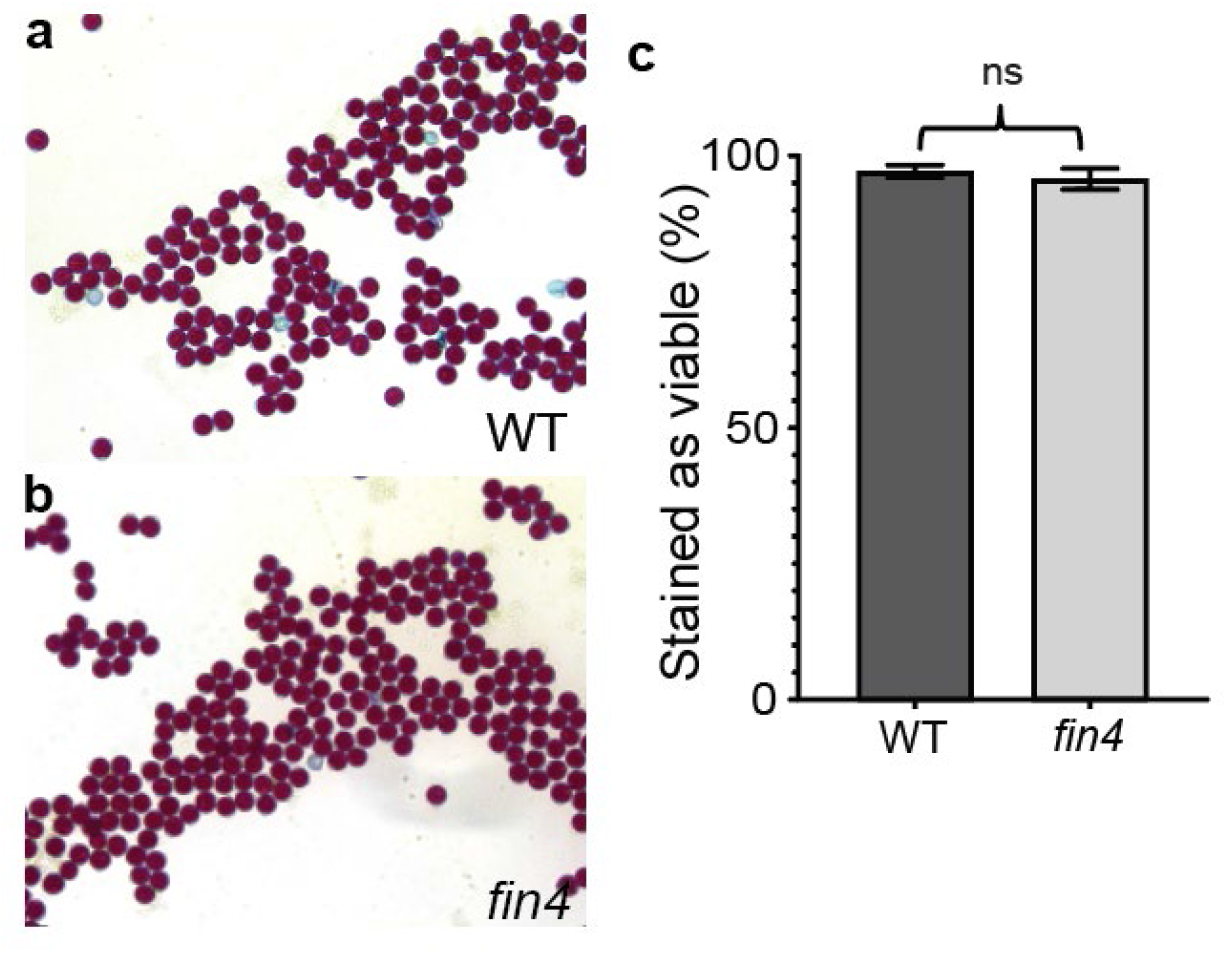
*fin4* pollen is viable. Alexander staining of WT (**a**) and *fin4* (**b**) pollen showing no difference in viability between the pollen of the two genotypes. Scale bar = 50 μm. **c.** Quantification of pollen viability as measured by Alexander staining of WT and *fin4* pollen.

**Supplemental Figure 4.**
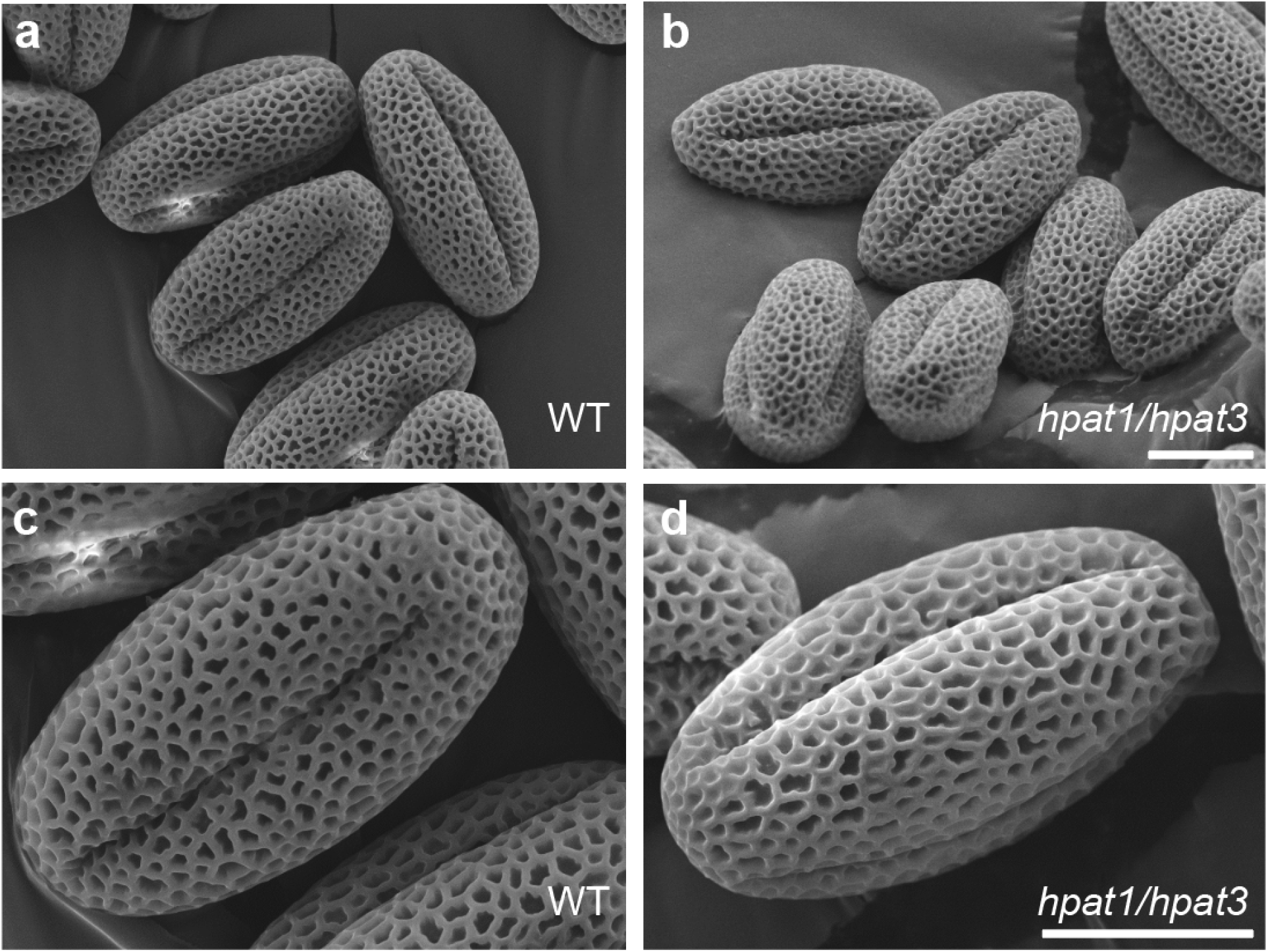
Arabidopsis *hpat1 hpat3* pollen do not collapse under SEM sample preparation conditions. SEM of Arabidopsis pollen grains from Columbia-0 wild-type (**a** and **c**) and *hpat1 hpat3* pollen (**b** and **d**). Scale bars = 10μm; a and b are at the same magnification as are c and d.

## Notes

### Competing Interest Statement

The authors have declared no competing interest.

